# Long-term mark-recapture and growth data for large-sized migratory brown trout (*Salmo trutta*) from Lake Mjøsa, Norway

**DOI:** 10.1101/544825

**Authors:** S. Jannicke Moe, Chloé R. Nater, Atle Rustadbakken, L. Asbjørn Vøllestad, Espen Lund, Tore Qvenild, Ola Hegge, Per Aass

## Abstract

**Background:** Long-term data from marked animals provide a wealth of opportunities for studies with high relevance to both basic ecological understanding and successful management in a changing world. The key strength of such data is that they allow to quantify individual variation in vital rates (e.g. survival, growth, reproduction) and then link it mechanistically to dynamics at the population level. However, maintaining the collection of individual-based data over long time periods comes with large logistic efforts and costs, and studies spanning over decades are therefore rare. This is the case particularly for migratory aquatic species, many of which are in decline despite their high ecological, cultural, and economical value.

**New information:** This paper describes two unique publicly available time series of individual-based data originating from a 51-year mark-recapture study of a land-locked population of large-sized migratory brown trout (*Salmo trutta*) in Norway: the Hunder trout. In the period 1966-2015, nearly 14,000 adult Hunder trout have been captured and individually marked during their spawning migration from Lake Mjøsa to the river Gubrandsdalslågen. Almost a third of those individuals were later recaptured alive during a later spawning run and/or captured by fishermen and reported dead or alive. This has resulted in the first data series: a mark-recapture-recovery dataset spanning half a century and more than 18,000 capture records. The second data series consists of additional data on juvenile and adult growth and life-history schedules from half of the marked individuals, obtained by means of scale sample analysis. The two datasets offer a rare long-term perspective on individuals and population dynamics and provide unique opportunities to gain insights into questions surrounding management, conservation, and restoration of migratory salmonid populations and freshwater ecosystems.

## Introduction

Important processes in ecology and evolution of vertebrates occur over the course of multiple years and often decades. Many areas of ecological and evolutionary research – including most of studies with the goals of improving species management and conservation – therefore rely on the availability of data spanning long time periods. Long-term ecological data on animal populations may be collected either at the population level (*e.g.* count or occupancy surveys) or by following individuals with uniquely identifiable marks throughout their lives. Individual-based mark-recapture and life-history data resulting from the latter provide a wealth of opportunities for studies that are impossible with only population-level data, as they not only allow linking population dynamics to vital rates (e.g. survival, growth, reproduction) but also enable to study individual differences in those vital rates (*Clutton-Brock and Sheldon 2010*). When mark-recapture and life-history studies are run over long time periods and include large numbers of individuals, as well as multiple cohorts and generations, study opportunities and investigable research questions multiply (*Clutton-Brock and Sheldon 2010*). However, maintaining the collection of individual-based data over long time periods comes with large logistic efforts and costs, and studies spanning many years are thus rare.

Recent declines of freshwater species abundance is more severe than species declines on land or in the ocean, according to the latest Living Planet report (*WWF 2018*): populations of freshwater species have declined by more than 80% on average during 50 years, while populations of land-dwelling- and oceanic species have fallen by less than 40%. Salmonid fishes, which are top predators and keystone species in many large freshwater ecosystems, are of high ecological, cultural, and economical value (*Lobón-Cerviá and Sanz 2018*). In spite of extensive study, however, many native wild populations remain in decline (*Muhlfeld et al. 2018*, *Muhlfeld et al. 2019*). Conservation concerns are particularly great for migratory salmonids as local adaptations and long life spans make them very vulnerable to environmental changes and habitat alteration, e.g. due to hydroelectric power production (*Piccolo et al. 2012*, *Van Leeuwen et al. 2018*). Habitat destruction and overexploitation from fishing are among the anthropogenic pressures threatening viability of salmonid populations (*Post 2013*).

Intense stocking with hatchery-reared fish, which can be necessary for sustaining a viable fish population, can nevertheless compromise the genetic integrity of wild populations (*Laikre et al. 2010*). More recently, climate change has emerged as an additional threat to populations of migratory salmonids globally (*Kovach et al. 2016*) but lack of knowledge and long-term data severely hamper the effectiveness of management and conservation efforts.

This paper describes two unique individual-based datasets that have resulted from a long-term study on a land-locked population of large-sized, piscivorous brown trout (*Salmo trutta*) in Norway, commonly referred to as Hunder trout (*Fig. 1*). The first dataset is a 51-year time series of mark-recapture-recovery data encompassing 13,975 adult trout captured and marked during spawning migration between 1966 and 2015. The second dataset consists of back-calculated age, growth and life-history information inferred from scales sampled from 6,875 of those marked trout in the period 1966-2003. These datasets offer rare opportunities to quantify vital rates and life-history processes that are central to salmonid life cycles such as somatic growth, developmental schedules, natural and fishing mortality, spawning biology, and the usage and effectivity of a fish ladder (*Nater et al. 2018*, *Nater et al. 2019b*, *Haugen et al. 2008*). Analyses building on both datasets can be used to gain insights into a variety of questions regarding management and conservation of migratory salmonid populations. These include – but are not limited to – consequences of recreational fishing, hydroelectric power production, stocking programmes, and environmental disturbance including habitat alteration and climate change.

**Figure 1.**
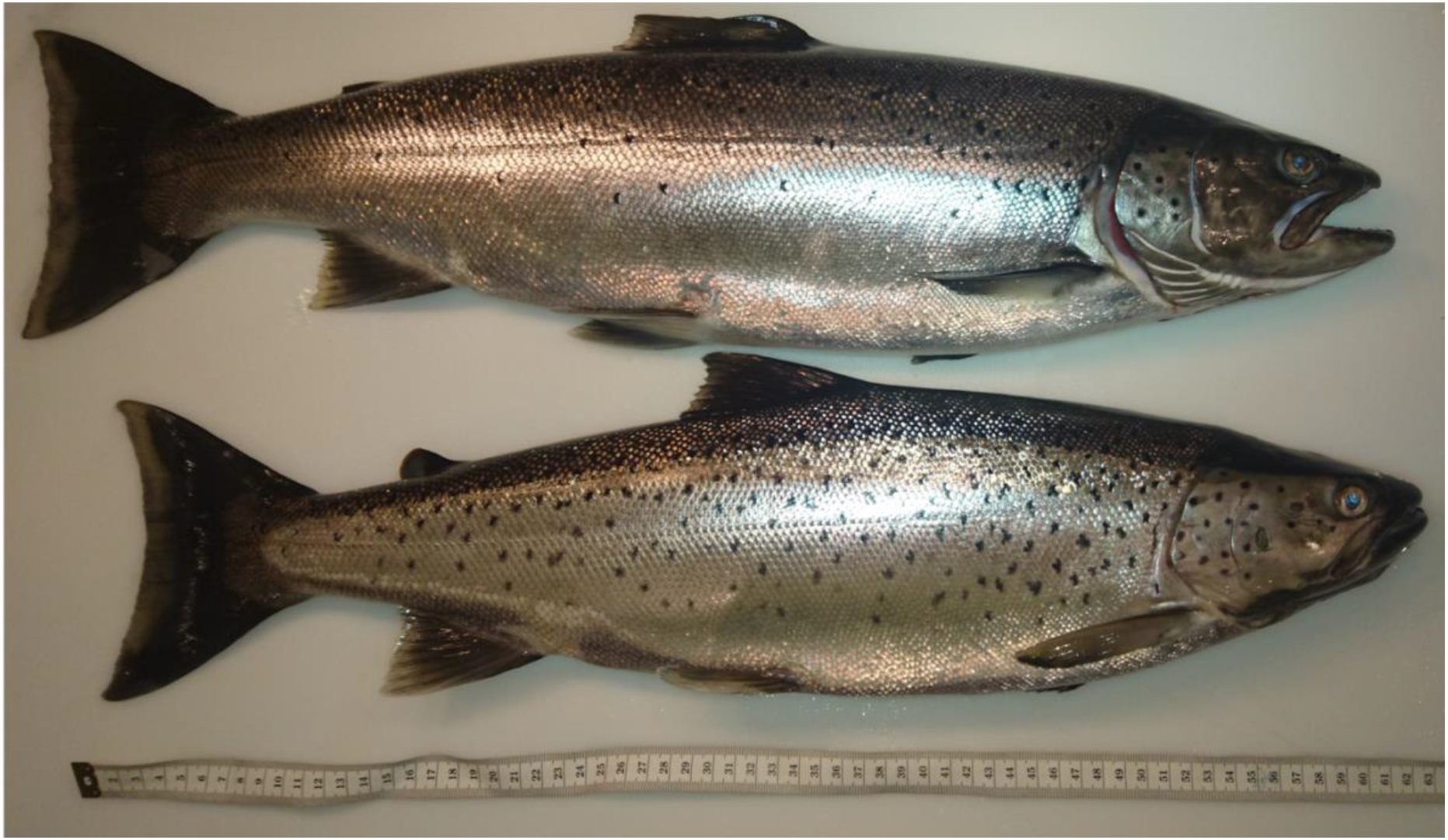
Two adult Hunder trout with body lengths exceeding 60 cm. The upper individual has a clipped adipose fin, indicating hatchery origin. Photo: Atle Rustadbakken.

## General description

We here describe a long-term study of the Hunder trout and provide documentation for the resulting (capture-)mark-recapture-recovery (CMRR) data and growth data with the aim of facilitating re-use of this unique data resource. We include important information on biology, habitat, study design and sampling protocols alongside detailed metadata for both datasets and notes on their explicit and implicit connections and potential uses.

### The Hunder trout population

The Hunder trout is a migratory brown trout inhabiting Lake Mjøsa and its main inlet River Gudbrandsdalslågen (also called Lågen) in Southeast Norway. The population is famous for its large body size (with some individuals measuring > 1 m and weighing > 10 kg), and its life history and spawning biology have been well studied (*Aass et al. 1989*, *Kraabøl 2006*, *Nater et al. 2018*). Although the population is land-locked (freshwater exclusive), its appearance and life cycle are very similar to that of the closely related anadromous brown trout (sea trout) and Atlantic salmon *(Salmo salar)*. Eggs are deposited in the river in fall and overwinter in loose gravel before hatching in the subsequent spring. Juvenile Hunder trout typically spend the first three to five years of their lives in the river while feeding on invertebrates. Upon reaching an average size of 25 cm, the young trout smolt and migrate downstream into Lake Mjøsa during the spring (corresponding to the seaward migration of anadromous trout and salmon). In the lake they shift to a piscivorous diet and this ontogenetic niche-shift is accompanied by an initially drastic increase in annual growth rate. Typically after another two to four years of rapid growth in the lake, the trout mature and migrate back to the river to spawn during the fall. At the time of their first spawning run, trout are on average 7 years old, weigh 3.5 kg and measure 63 cm. Following the first spawning run, the trout return to the lake and subsequently perform spawning runs every other year. The majority of individuals strictly adhere to this biennial spawning cycle, but cases of trout spawning in two or more subsequent years have been recorded (<1.5% of the population based on the here described long-term data). Prior to the damming of the Gudbrandsdalslågen (see below) a large part of the population is believed to have spawned upstream of the Hunderfossen waterfalls. Overcoming this natural obstacle during the spawning migration has been proposed as one of the drivers of selection towards the large body size of this population (*Haugen et al. 2008*) relative to that of trout populations spawning in other rivers draining into Lake Mjøsa (e.g. *Linløkken et al. 2014, Rustadbakken et al. 2004, Skaala 1992*).

### Anthropogenic impacts: harvest, hydropower, and mitigation measures

The Hunder trout has a long history of being impacted by human activity both directly through harvest and indirectly through habitat alteration.

The Hunder trout has been the target of intense harvest for centuries; see *Aass and Kraabøl 1999*for an overview of the history of the trout fishery. Fishing mortality is a key factor determining adult survival (*Nater et al. 2019b*) and driving population dynamics (*Nater et al. 2019a*a). Over the course of the study period (1966-2017), fishing was done mainly by means of gillnets and angling and its objective shifted gradually from subsistence to recreation. Catch-and-release practices were very rare during the study period, but have gained popularity during the last decade.

Two major dams were established in River Gudbrandsdalslågen in the 1960s, the Harpefoss and Hunderfossen dams (Figure 2). The latter dam, built between 1961 and 1964, is located in the historical spawning grounds of the Hunder trout and thus constitutes a migration barrier. To alleviate the impacts of this barrier and restore the connectivity of the river, a fish ladder was established within the Hunderfossen dam in 1966 (*Aass 1993*). The functionality of this fish ladder and its success in allowing trout to pass the dam on their upriver migration vary among seasons and depend heavily on water temperature and flow, as well as body size of the individual trout (*Jensen and Aass 1995*, *Haugen et al. 2008*, *Nater et al. 2019b*, *Kraabøl and Museth 2019*. This fish ladder is in practice a one-way passage and facilitates only upriver migration. Consequently, the dam still poses a major obstacle for downriver migrating smolt and post-spawners. Mortality associated with passage via floodgates or turbine shaft, increased predation in the dammed area and migration delays are thus likely to affect trout survival during the downriver passage (e.g. *Fjeldstad et al. 2018*).

**Figure 2.**
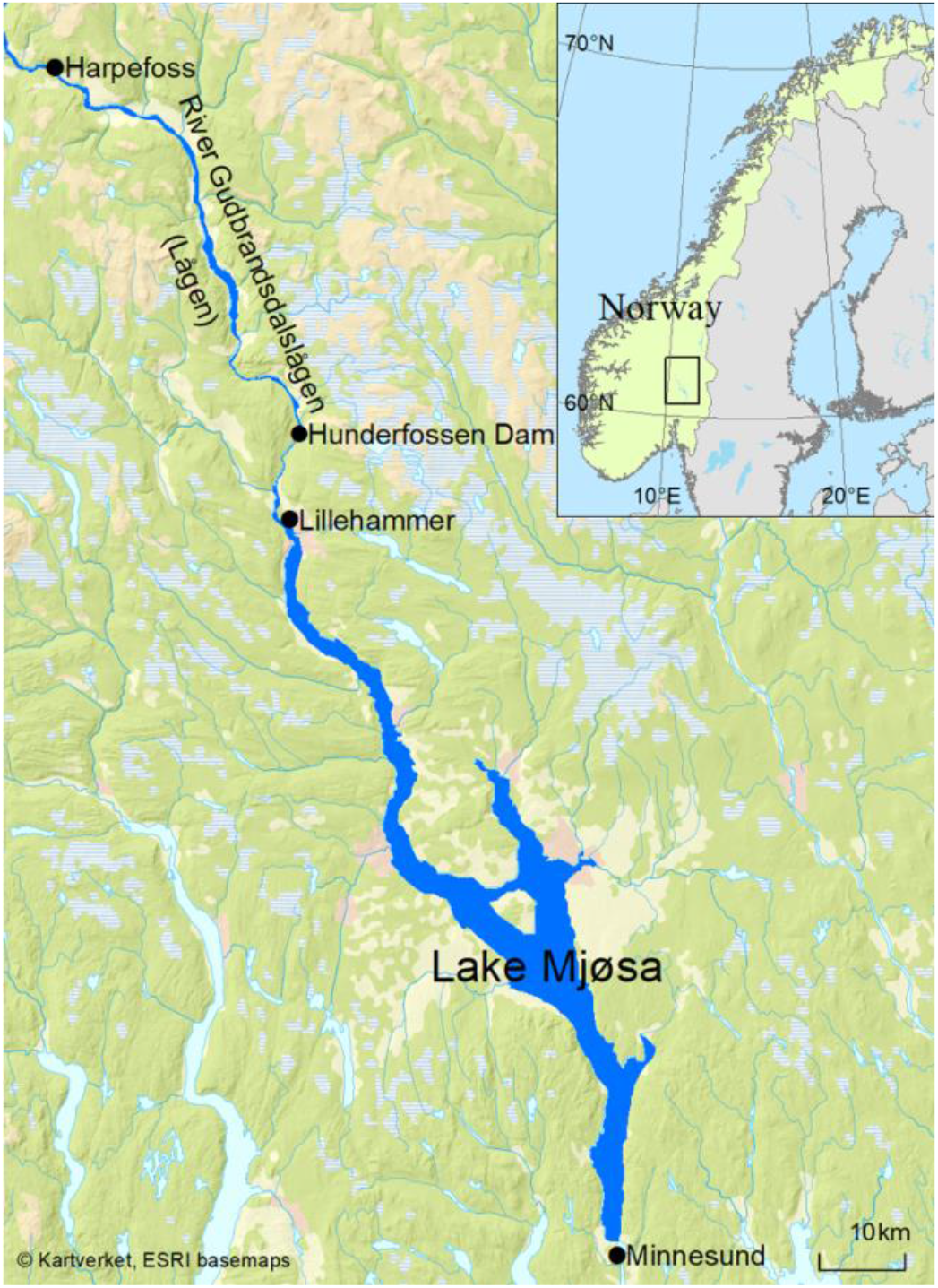
Map of Lake Mjøsa and the river Gudbransdalslågen, where the Hunder brown trout occurs. The fish ladder and the hatchery are located at the Hunderfossen dam. Sources: The Norwegian Mapping Authority (https://www.kartverket.no) and ESRI (https://www.esri.com). After *Aass et al. (2017)*.

Damming also drastically altered the hydrological conditions in the river, leading to a reduction in availability and quality of spawning and nursing habitats in the river both upstream and downstream of the dam (*Aass et al. 1989*). To compensate for the resulting decreased natural production of trout in the regulated river, a stocking programme was initiated in the mid-1960s (*Aass 1993*). This programme entails the release of between 10,000 and 30,000 first-generation hatchery-reared smolts into the river and lake each year and is still being continued as of today. Every year, eggs and sperm are collected from a fraction of the wild-born spawners that ascended the fish ladder. These are incubated in a hatchery next to the Hunderfossen dam, and the hatchling trout are reared until smolting (usually for 2 years) before being stocked into the population (with their adipose fins clipped to allow recognition later).

## Project description

### Project

Sustainable management of renewable resources in a changing environment: an integrated approach across ecosystems (SUSTAIN). https://www.sustain.uio.no/

### Work Package 1

Demographic structure in harvested ecosystems. https://www.sustain.uio.no/research/wp/work-package-1/

### Funding

Research Council of Norway, project SUSTAIN, contract no. 244647/E10. https://www.sustain.uio.no/

## Study area description

### Lake Mjøsa and River Gudbrandsdalslågen

Lake Mjøsa is a deep fjord lake (max. depth 453 m) situated in Southeast Norway (*Fig. 2*). The surface area is 366 km2, mean depth is 155 m, and the residence time is 5.6 years. It has a large catchment of 16.6 km2 with approximately 200,000 human inhabitants living close to the lake. The Southern parts of the catchment are dominated by pine forests, while the northern parts lie within mountainous regions, and several glaciers in this region feed the main tributary rivers with heavy silt load resulting from snow-melt during June-August. The water flow from the main river, Gudbrandsdalslågen, as well as several smaller inlets, causes reductions in transparency, temperature and algal growth in Lake Mjøsa during early summer, especially in the Northern parts of the lake (*Holtan 1979*). More details on the pressures, ecological status and other properties of Lake Mjøsa and its catchment are available at https://vann-nett.no/portal/#/waterbody/002-118-1-L.

In the 1970s and 1980s, eutrophication due to excessive nutrient loads from agriculture, industry and households resulted in poor water quality and harmful algal blooms in Mjøsa. A subsequent large restoration effort (the Lake Mjøsa campaign) resulted in a period of re-oligotrophication and the current good ecological status of the lake (*Lyche Solheim et al. 2019*). The campaign also marked the start of extensive monitoring of water quality and plankton communities in Lake Mjøsa, the outcome of which are more than 40 years of physico-chemical and biological time-series data (*Løvik and Moe 2016*). In addition to the information on changes in water quality, these data have also revealed a trend of increasing water temperature (*Hobæk et al. 2012*), as well as shorter ice coverage periods and an increased frequency of floods in more recent years.

## Sampling methods

The establishment of the fish ladder within the Hunderfossen dam resulted in unique opportunities for monitoring the spawning population of the Hunder trout. Concurrent with the opening of the fish ladder in 1966, a mark-recapture programme was initiated and was run continuously until termination in 2016. This monitoring program has resulted in two unique long-term data series: a 51-year time series of (capture-)mark-recapture-recovery (CMRR) data and a 37 years of scale samples resulting in a 51-year time series of growth data (*Fig. 3*).

**Figure 3.**
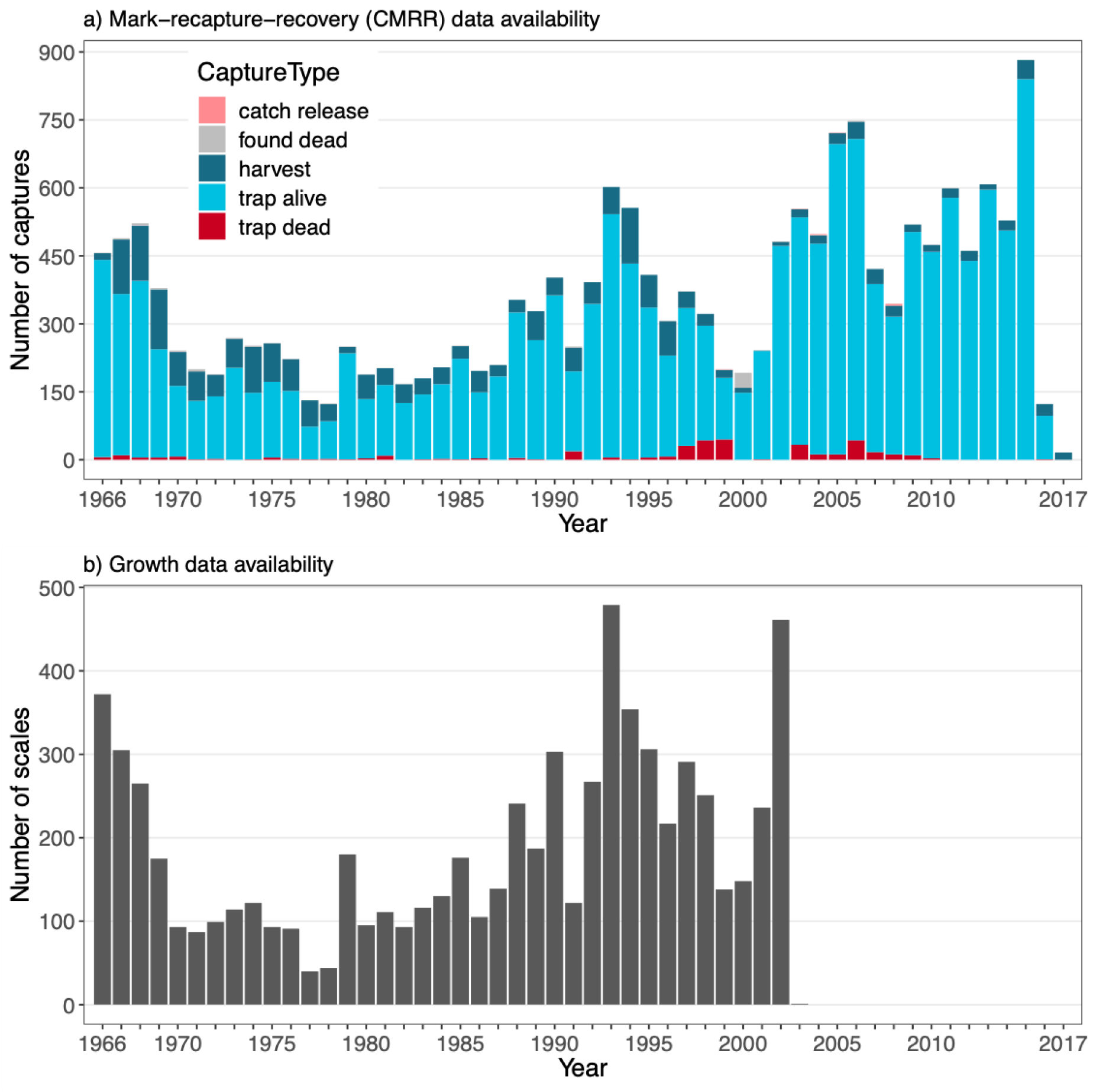
Overview of availability of (a) (capture-)mark-recapture-recovery (CMRR) data and (b) growth data over the course of the study period (1966 - 2017). For CMRR data, counts of trout captures are categorised by capture type (see colour legend). Captures of types “trap dead” and “trap alive” occur in the trap within the Hunderfossen fish ladder; “harvest” and “catch release” are captures by fishers in any location and resulting in death of the fish for the former and alive re-release for the latter; capture type “found dead” is assigned to fish that are discovered already dead in any location except for the trap within the Hunderfossen fish ladder. For more details, see Table S.2 in *Suppl. material 1*.

In the following, we describe the generation of mark-recapture data from trapping of spawning trout in the Hunderfossen fish ladder, its supplementation with recovery data obtained as reports of marked fish by local fishers, and sampling and sclerochronological analysis of scales resulting in data on individual growth and life-history scedules.

All data were originally collected, recorded, maintained, and processed by a variety of individuals and institutions, as listed in the Author contributions and Acknowledgements sections.

### Marking and recaptures in the fish ladder

All adult trout ascending the Hunderfossen hydropower dam on their spawning migration between 1966 and 2016 were captured in a trap situated in the lower part of the fish ladder. Until 2015, all unmarked trout were individually marked with a Carlin tags, consisting of a plastic disc with a unique mark number (alphanumeric code) and a stainless steel thread to attach to the fish (*Fig. 4*, *Carlin 1955*). Trout that already had a Carlin tag from a previous capture had their mark number redistered. All trapped fish were measured by measuring tape (precision 1 cm) and weighed with a Salter Brecknell scale (precision 100 g). Furthermore, each individual’s sex was determined if possible based on secondary sex characteristics such as colour, presence of a kype and sexual products (eggs and sperm). Its origin (wild-born or stocked) was ascertained by checking whether the adipose fin was intact (removed = stocked). All handling were done under anaesthesia. After registration and sampling, trout were released into a resting pool connected to the second part of the fish ladder, allowing them recover and then complete their dam passage and migration to the upriver spawning areas. If an individual was unable to recover and died under sampling or in the resting pool, this was also recorded.

**Figure 4.**
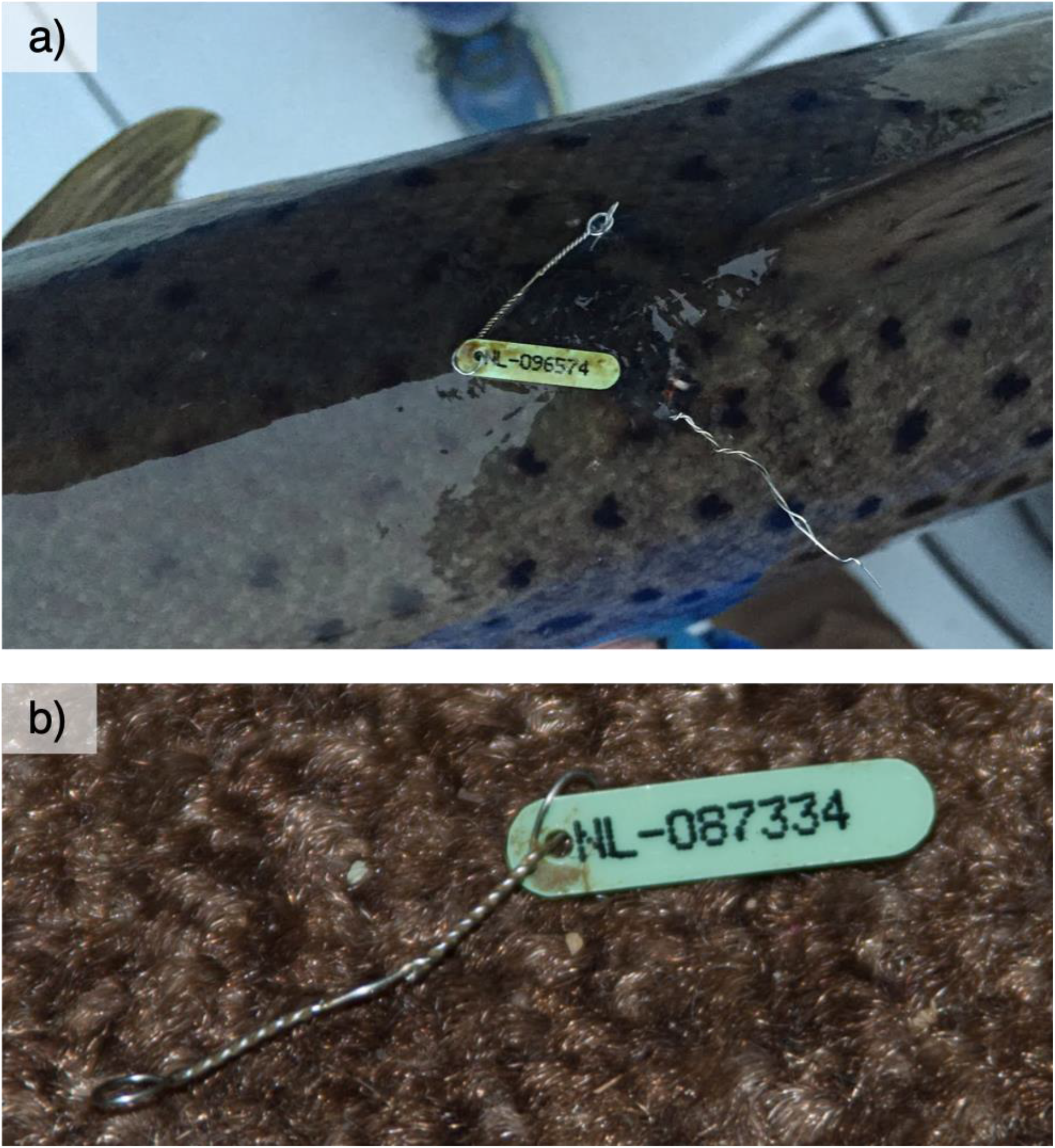
(a) Fish marked with a Carlin tag. Photo: Frank Ronny Johansen. (b) Detail of a Carlin tag with individual code. Photo: Atle Rustadbakken.

### Recoveries from fishing reports and others

Marked trout could be recaptured in the fish ladder during later spawning runs as well as by fishers in any part of the river or lake. Reports of such harvest recoveries are available from 1966 to 2017. When trout were caught by fishers and reported, the amount of data collected varied. As a minimum, fishers reported the mark number of the caught fish, the date of capture, and whether the fish was killed or released. Additional information could include capture location, fishing gear, sex, origin, length, and weight. Reporting the capture/harvest of a marked trout was done on a voluntary basis, and no monetary reward was given in exchange for reports. In combination with declining feedback to fishers from the marking project, this likely led to a gradual decrease in fisher’s reporting rate over the course of the study period (*Nater et al. 2019b*).

On rare occasions, marked trout were found dead and reported by people that did not capture them through fishing themselves. This includes, for example, trout that died while attempting dam passage and were found dead in the vicinity of the hydropower station, trout that died naturally and washed up on the shores of the lake or the river, and even reports from people which had purchased a marked trout on a fish market. All of these reports were included in the CMRR dataset.

### Scale sampling and sclerochronological analysis

During the period 1966-2015, 4-6 scales were obtained from a large number of trout captured in the fish ladder. The scales were sampled from a standardized location above the lateral line between the dorsal and the adipose fin using small forceps. For a subset of individuals that ascended the fish ladder between 1966 and 2003, the sampled scales have been analysed using schlerochronological methods (*Bagenal 1978, Panfili et al. 2002*). Sclerochronological analyses allow to backcalculate fish size and age from growth increments in the scales (*Fig. 5*) under the assumption of an isometric relationship (Lea-Dahl method, *Bagenal 1978*). Additionally, analysis of sclerite patterns (growth righs deposited in the scale during somatic growth) may allow the determination of the presence, timing, and frequency of key life-history events. For the Hunder trout, the smolting event (mark “S” in *Fig. 5*) is indicated by the transition from relatively slow river growth (denser sclerites) to faster lake growth (more space between sclerites). Spawning events are recognizable by the typical spawning marks they leave on the scale, including erosion of outer parts of the scale, broken sclerites, new sclerites crossing over older ones. Drastic reduction in growth is a typical consequence of energy allocation from somatic to gonad growth before and during spawning runs. Sclerochronolocial analysis of Hunder trout scales thus provided important life-history information: juvenile growth in the river, sub- and post-adult growth in the lake, age and size at smolting and maturation and the number of previous spawning events and resting years.

**Figure 5.**
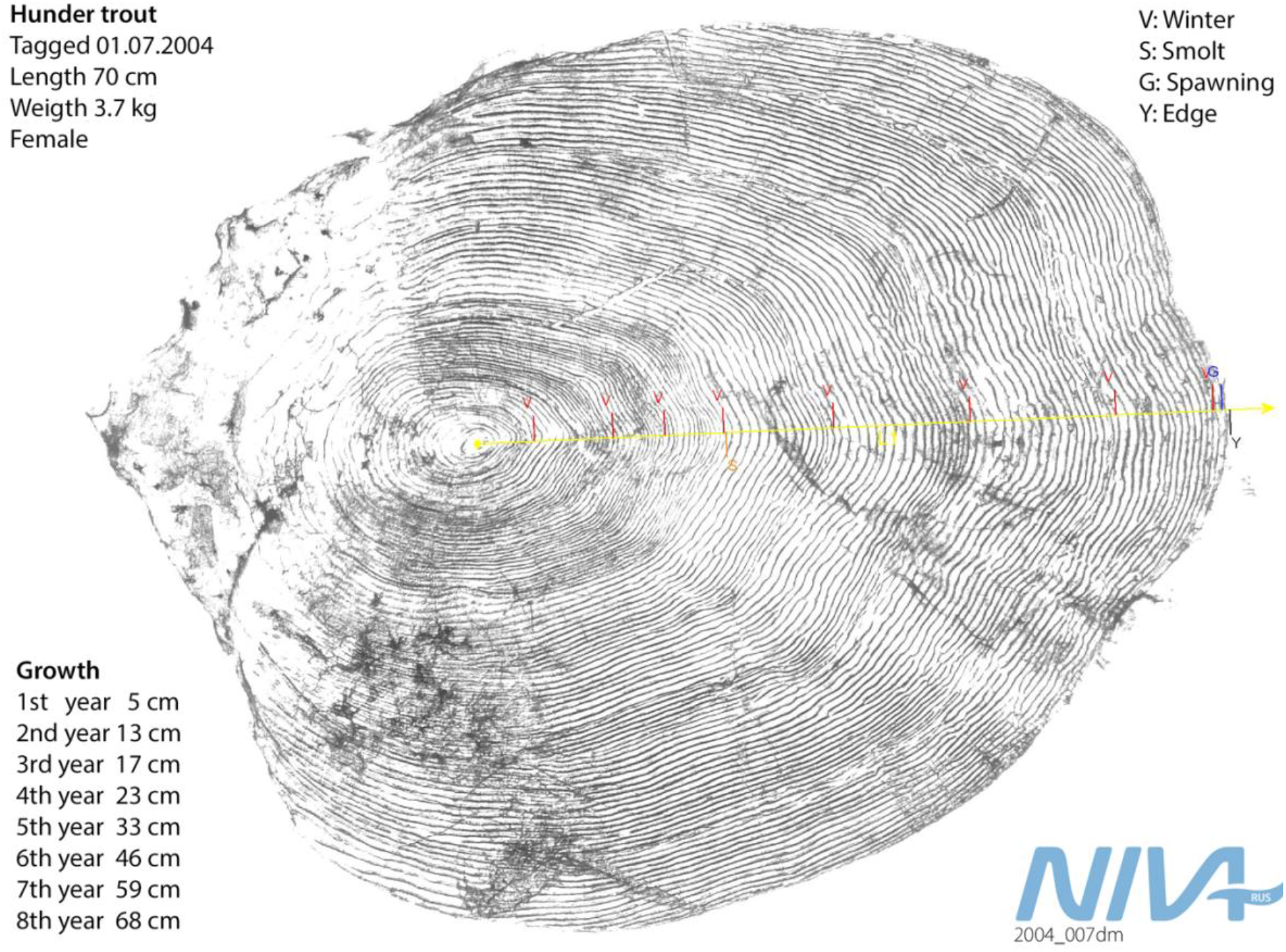
Illustration of a fish scale and interpretation of growth, development and spawning events. The yellow line represents the time line from hatching (centre) to the current age (the edge). Red lines indicate winters (marked “V”), where the growth is slower (denser sclerite pattern). Smoltification (“S”) after the 4th winter is indicated by subsequent rapid growth. Spawning runs (“G”) are followed by very slow growth during the summer propor to and during the spawning run.

## Geographic coverage

### Description

The spatial extent of the river and lake system the Hunder trout inhabits spans the latitudes 60.40° - 61.22°N and longitudes 10.43° - 11.29°E (GCS, WGS84) (*Fig. 2*). The altitude ranges from 123 m a.s.l. (lake surface) to 175 m a.s.l. (the upper end of river spawning areas).

### Coordinates

60.40 and 61.22 Latitude; 10.43 and 11.29 Longitude.

## Taxonomic coverage

### Description

The data are from a single population of large-sized migratory brown trout (*Salmo trutta*), also referred to as Hunder trout, inhabiting Lake Mjøsa and River Gudbrandsdalslågen.

### Taxa included

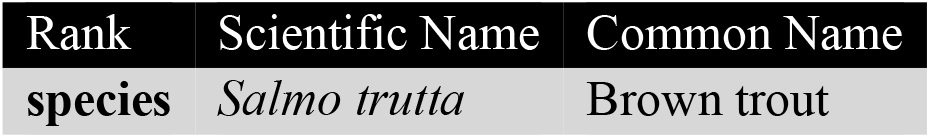

## Temporal coverage

The (capture-)mark-recapture-recovery (CMRR) data contains captures from all years in the period 1966-2017. As individual marking was discontinued after the 2015 sampling season, no new individuals were marked in 2016 (recaptures and recoveries only) and only harvest reports were recorded for 2017.

Scales were sampled in the period 1966-2015. Scales sampled during the years 1966-2003 have been analysed so far. Individual growth data is therefore available from 1952 (due to back-calculation) to 2003.

## Usage rights

### Use license

Creative Commons Public Domain Waiver (CC-Zero)

### IP rights notes

The data described here has been made publicly available. We do note, however, that successful re-use of long-term individual-based data requires good understanding of both data structure and the biological system, as well as an overview over previous work using the data, whether that work has been published in a scientific journal or not (see *Mills et al. 2015* for a perspective on opportunities and challenges associated with re-use of long-term ecological data).

For the above reason, we encourage parties interested in re-using the here described data to contact the authors CRN, SJM, AR or LAV for a discussion of research ideas and potential opportunities for collaboration.

## Data resources

**Data package title:** SUSTAIN trout data

**Resource link:** doi:10.5061/dryad.9cnp5hqf4 (not active yet)

**Number of data sets:** 2

### Mark-recapture-recovery data

**Data set name:** Hunder trout mark-recapture-recovery dataset

**Character set:** UTF-16

**Download URL:** TBA

**Data format:** comma-separated values (csv)

**Description:** The **mark-recapture-recovery dataset**(**CMRR**) contains altogether 18,488 capture records from years 1966-2017. Of these, 13,975 are individual marking events of mature trout in the ladder. The remaining 4,513 capture records are recaptures by different means (as defined by **CaptureType**): 2,106 recaptures in the trap (376 of which resulted in death); 2,322 harvest reports (1,944 in the lake, 358 in the river, 20 in other or unknown locations); 72 reports of fish found dead; and 9 reports of catch-and-release fishing. An overview of all recorded captures across years is given in *Fig. 3*a, while average monthly numbers of captures are plotted in *Fig. 6*. *Fig. 7* visualizes the ratio of captures of stocked versus wild indviduals and the ratio of male versus female individuals over time. The growth dataset, which represents a subset of the CMRR dataset, results in similar ratios as the larger dataset in both cases. Size distributions (at capture) of fish sampled in the fish ladder are shown in *Fig. 8*. For more information, see *Suppl. material 1*.

**Figure 6.**
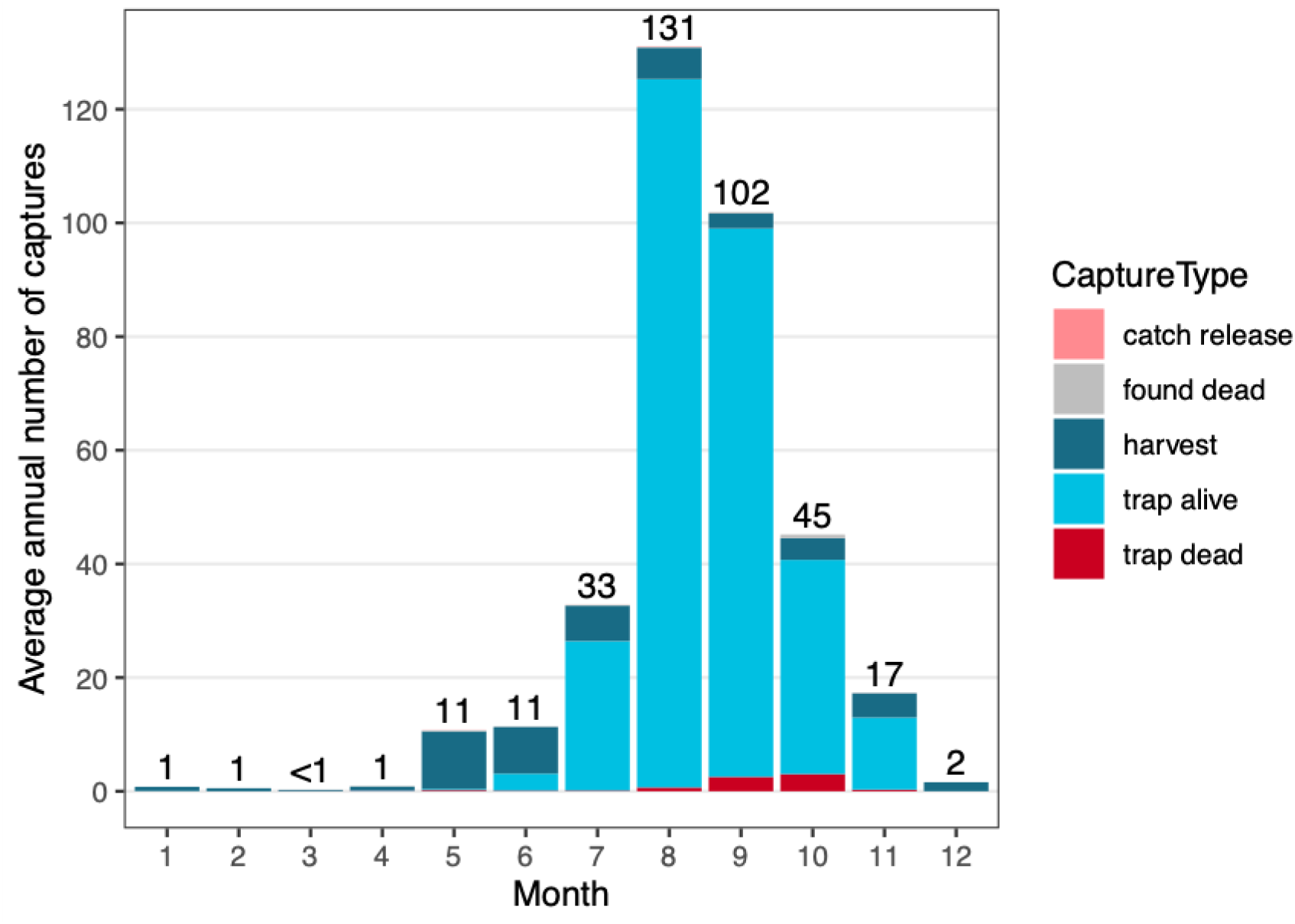
Distribution of trout captures across months for the period 1966-2016, categorised by capture type and averaged over all years. Digits above bars represent average numbers of all captures (all capture types combined) per month. For more details, see description of Figure 3.

**Figure 7.**
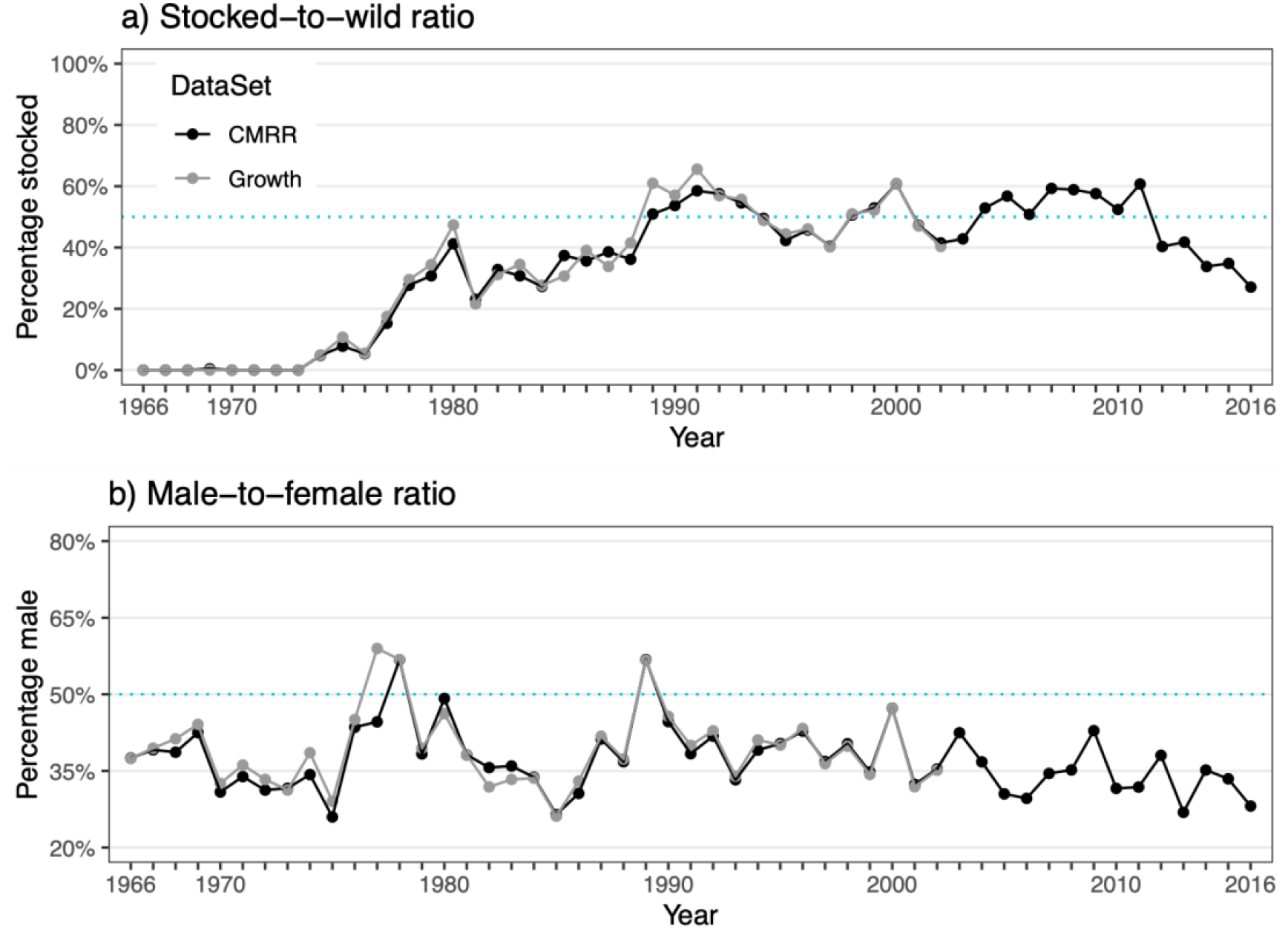
Ratios of (a) origin (stocked vs. wild) and (b) sex (male vs. female) of trout captured alive in the fish ladder (black) and scales sampled (grey) over the course of the study period. Ratios for scales samples in 2003 are not depicted, since only a single scale is available for this year. The dotted blue line marks the line of equal ratios (50%). Note that y-axis scaling differs for panels a) and b).

**Figure 8.**
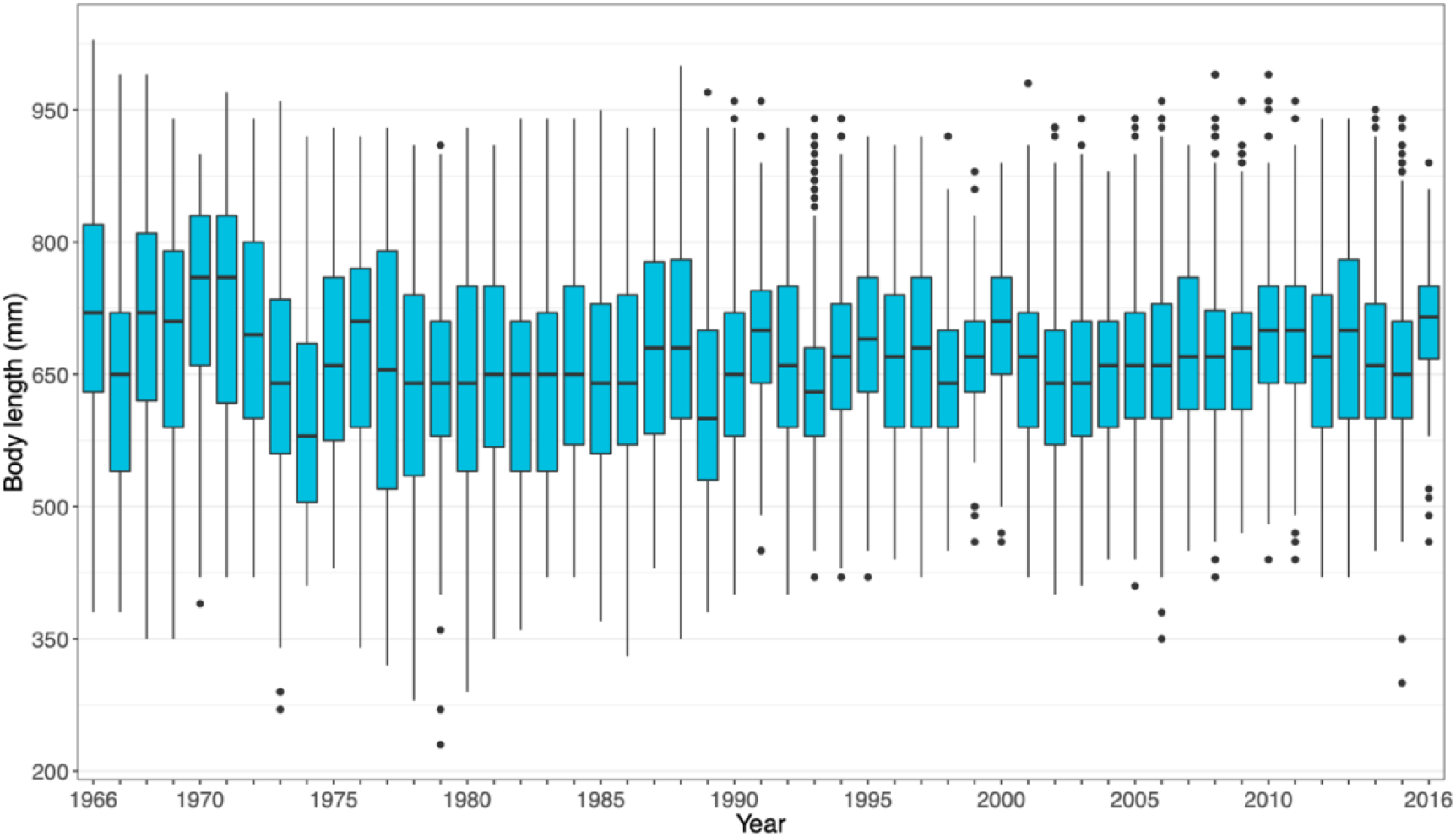
Annual distributions of individual body lengths measured upon capture in the fish ladder. Median body length is indicated by a white line. The box plot shows the median (horizontal bar), 25% and 75% quantiles (range of the box), largest and smallest values within 1.5 times the interquartile range (whiskers = vertical lines) and outliers (dots beyond whiskers).

**Metadata**:

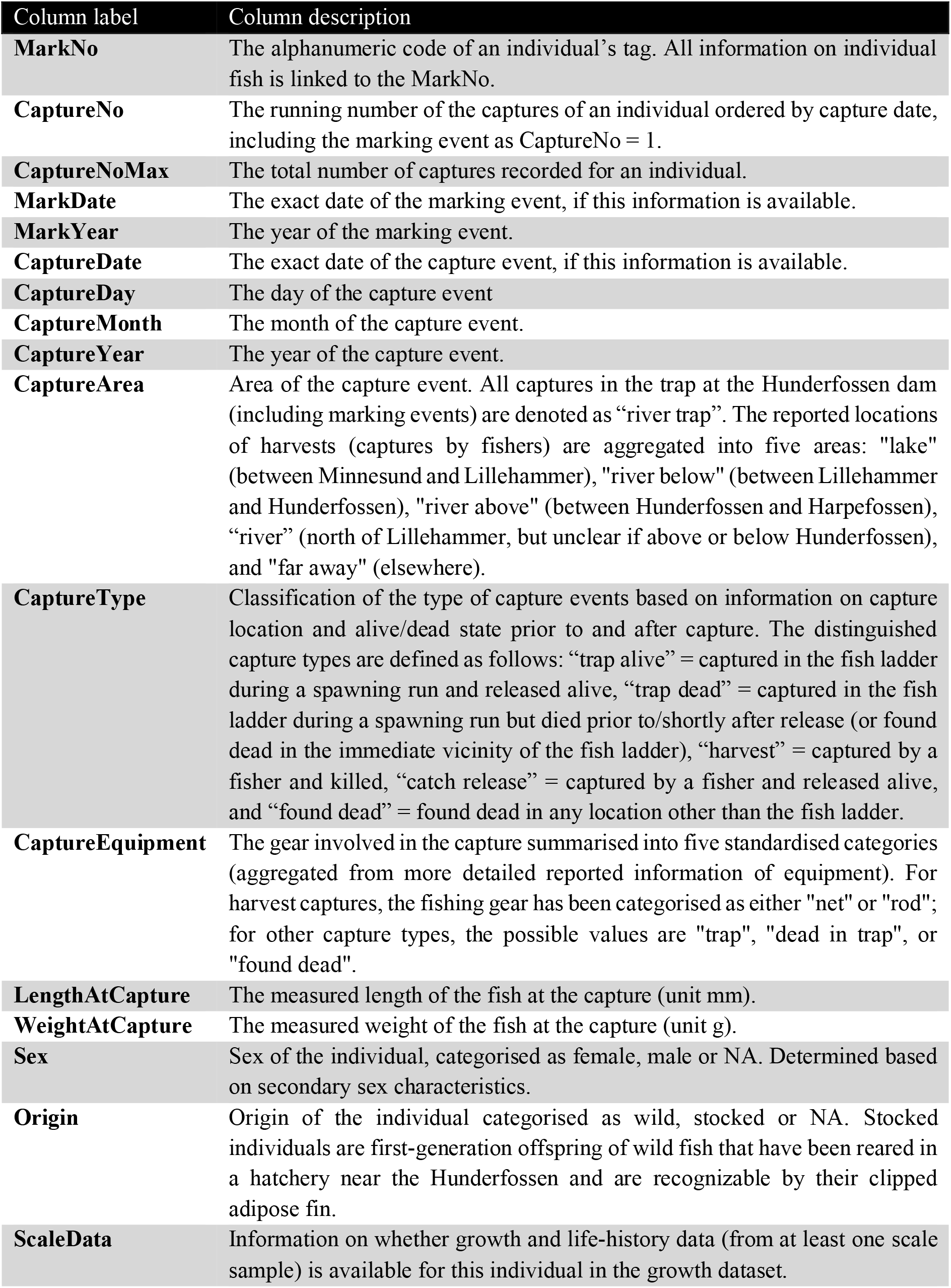

### Growth data

**Data set name:** Hunder trout growth dataset

**Character set:** UTF-16

**Data format:** comma-separated values (csv)

**Description:** The **growth dataset** contains altogether 41,605 size records back-calculated from 7047 scale samples collected between 1966 and 2003. The scales originate from 6,875 marked individuals, which are also present in the mark-recapture-recovery (CMRR) data. For the majority of individuals, growth data is available from a single scale sample in the dataset. 161, 10, and 1 individual(s) have growth data from two, three, and four scales collected at different capture events, respectively. Backcalculated sizes are divided into the juvenile period of life spent in the river and the adult period of life spent predominantly in the lake, and are supplemented with information on key life history events (smolting, maturation, spawning status). An overview of all sampled scale across years is given in *Fig. 3*b. *Fig. 7* visualizes the ratios of sampled scales from stocked versus wild and male versus female individuals over time. For more information, see *Suppl. material 1*.

**Metadata:**

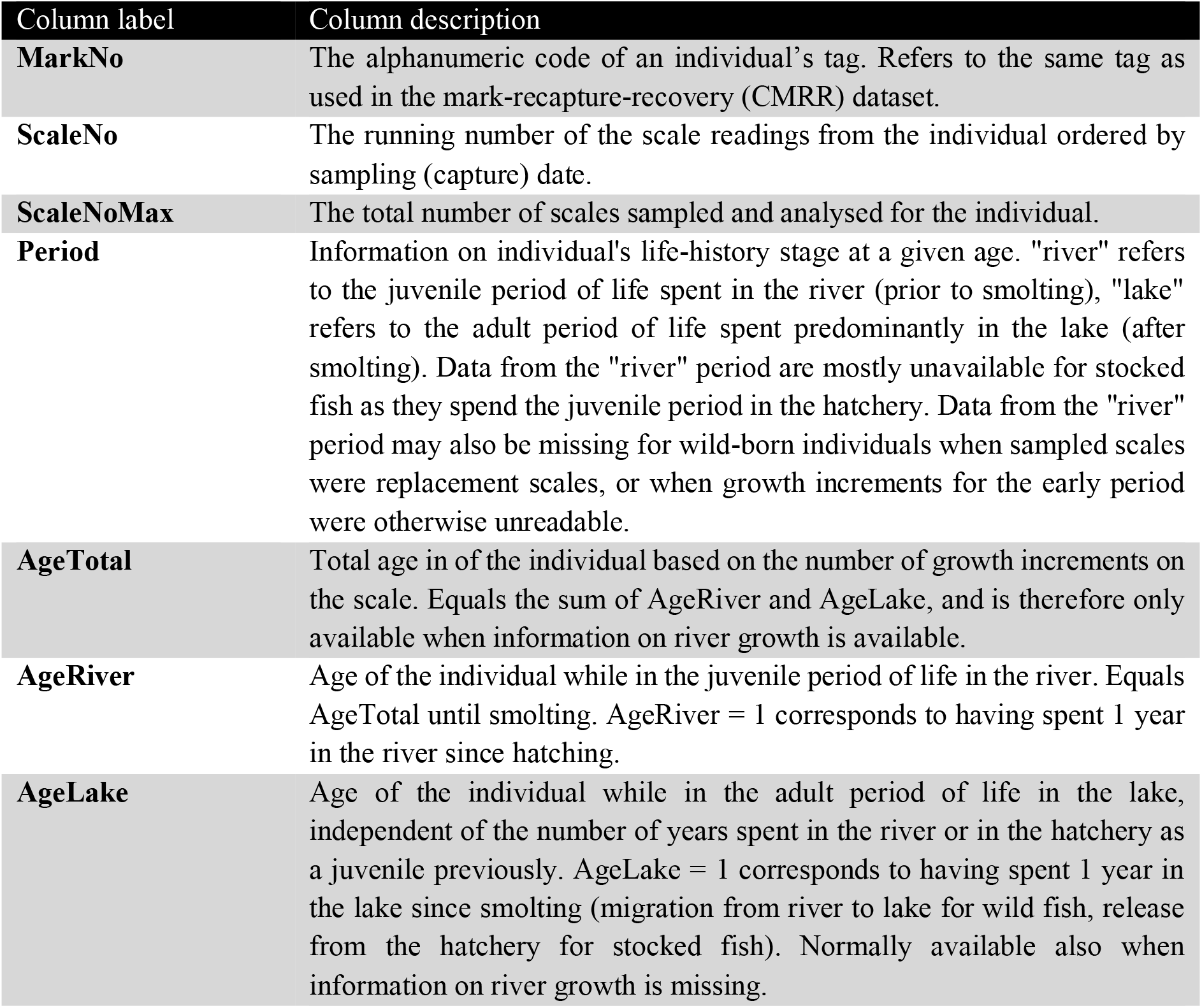
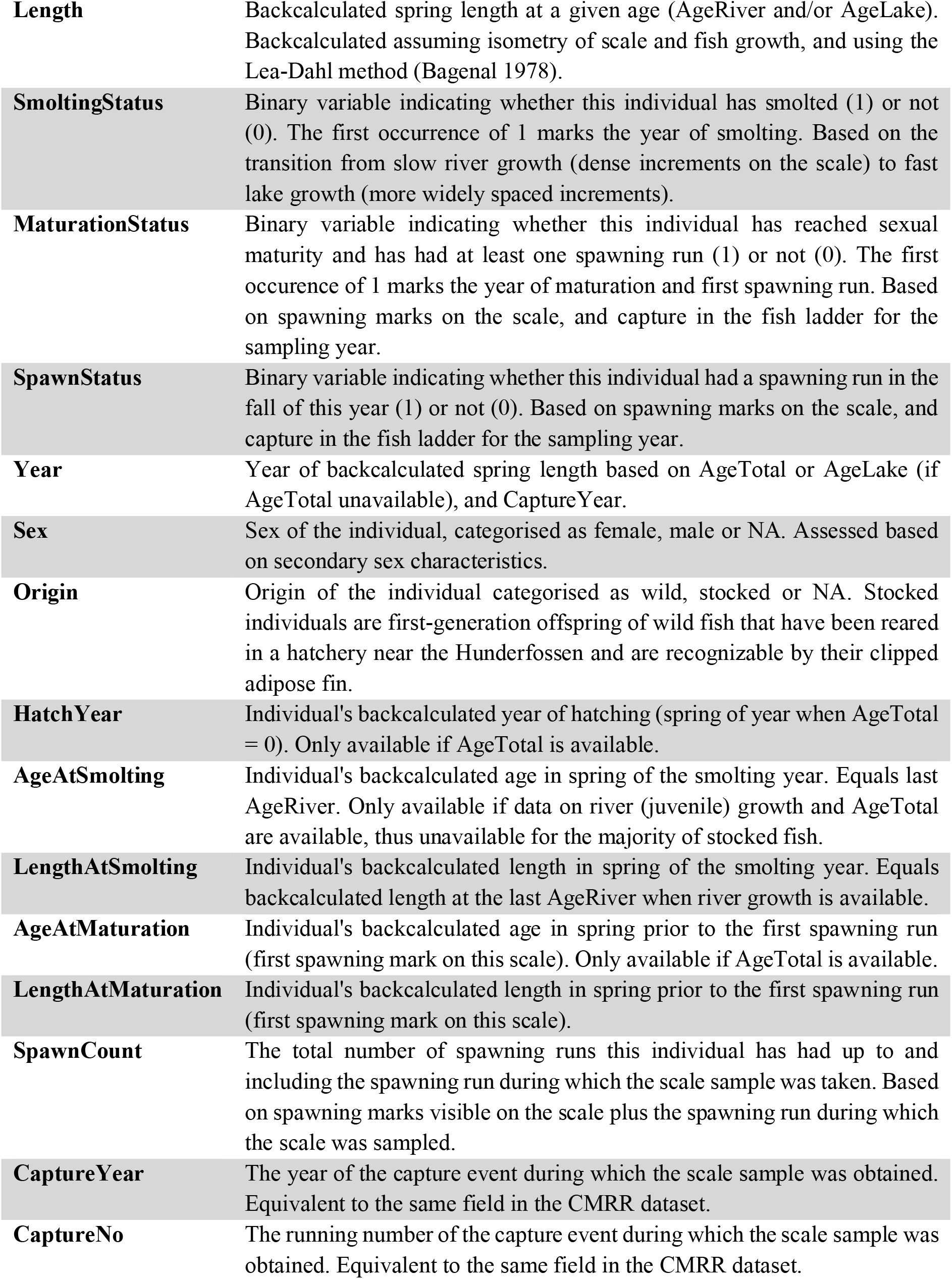

## Additional information

### The SUSTAIN trout database

The SUSTAIN trout database is a compilation of all available data on individuals, captures and scale-based information (growth and life history schedules) available for the Hunder trout, from which the two datasets described in this paper have been derived. For further use of the hunder trout data we recommend using the published datasets. To better explain the context and the data processing underlying the published dataset, we provide a more thorough description of this database and its structure in *Suppl. material 1*.

### Links between the CMRR and growth datasets

The above described CMRR and growth data are linked in the relational database as described in *Suppl. material 1*. The unique key linking the two datasets consists of the individual mark numbers (MarkNo) and the capture numbers (CaptureNo), which in combination defines unique capture/sampling events. All growth data originates from scales that were collected from marked trout ascending the fish ladder, meaning that for all individuals present in the growth data, mark-recapture-recovery data is also available. The opposite is not the case, since scale samples were only collected for a subset of all trout ascending the fish ladder, and not all collected scale samples were suitable for sclerochronological analysis.

An individual’s body length measured at capture in the fish ladder (LengthAtCapture) during the spawning season is used as the starting point for the backcalculation of size using a scale collected during the first capture. Since increments on scales are used to backcalculate spring size, however, this is done under the assumption that a trout’s length measured during the spawning run in the fall is roughly equal to its spring length earlier in the same year. This is not unreasonable, since growth in spawning years is greatly reduced relative to non-spawning years, and close to negligible for larger individuals (e.g. *Nater et al. 2018*).

Scale samples for some individuals have been obtained not (only) during the first capture in the fish ladder, but (also) during later recaptures in the fish ladder. For such individuals, information on body length in certain years may thus be available from both scale backcalculations and actual growth measurements during capture in the fish ladder. By assuming no growth during the winter (a reasonable assumption for brown trout, *Elliott 1994*), backcalculated spring size for any year t can be directly compared to measured fall size during a spawning run in year t-1. Such a comparison can be used to validate the sclerochronological analyses and show that lengths backcalculated from scales represent actual measured lengths well (*Fig. 9*, but see Appendix S2 in *Nater et al. 2018* for a more detailed treatment). Furthermore, the relationship between backcalculated spring lengths in years t and measured fall lengths in years t-1 allows for linking the two datasets in powerful integrated analyses.

**Figure 9.**
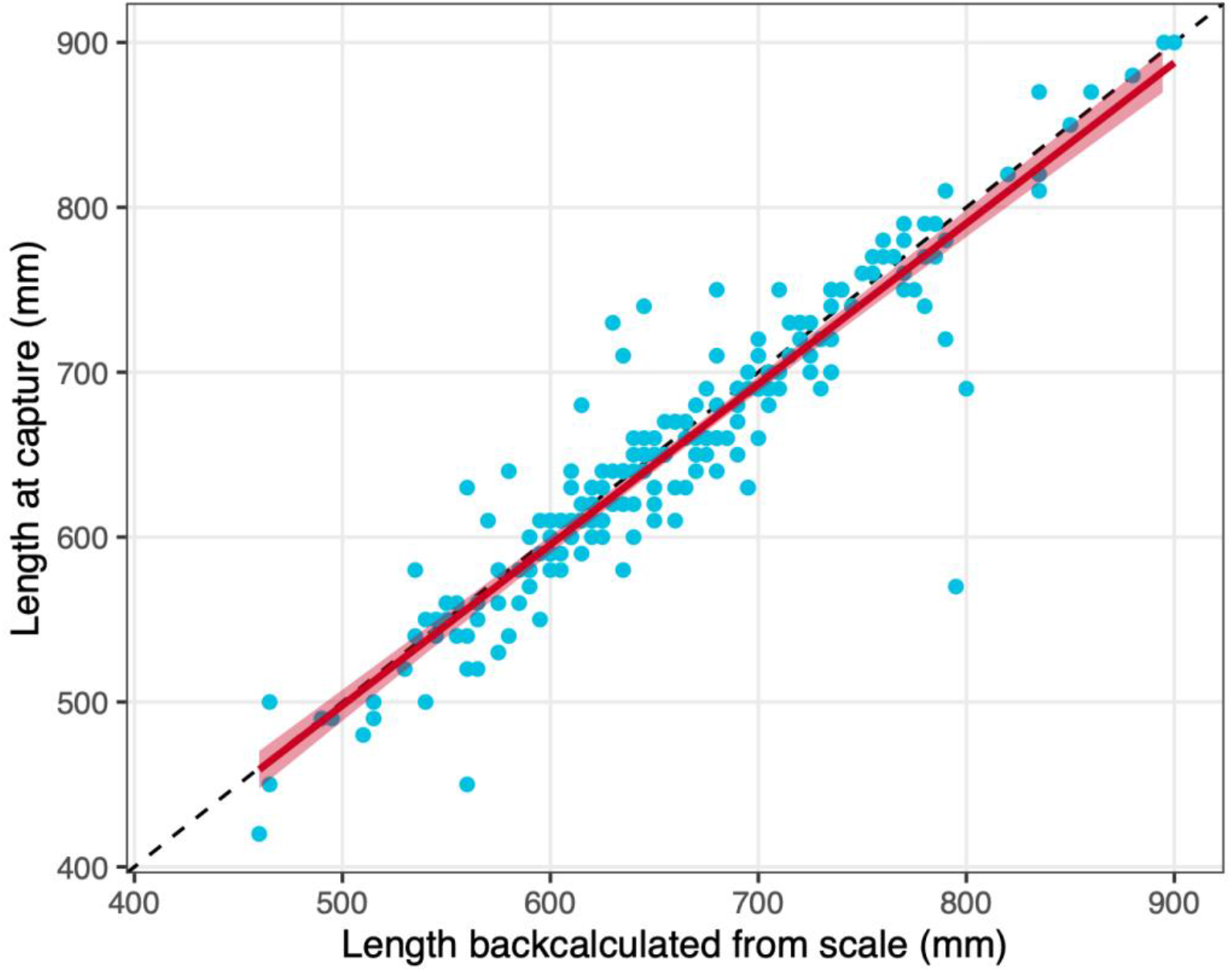
Correlation of backcalculated length from scales in spring (year t) and measured length in the fish ladder in fall the previous year (t-1). The blue dots represent data points (N = 201), the dashed line marks the 1-to-1 perfect correlation, and the red line is the fit of a linear regression including its confidence interval (intercept = 10.726, slope = 0.974, adj. R_2_ = 0.875).

### Availability of related Hunder trout data

Besides the two here described long-term individual-based datasets, additional data and samples have been collected on the Hunder trout in connection with the stocking programme and/or as part of shorter-term projects. Some of these data/samples may be made available on request by their respective owners, and we briefly mention some of them here due to their potential value for integration with the here described long-term data in future studies.

#### Additional scale samples

Scale samples were collected continually for the period 2004-2015 under the same protocol as described in this article. These additional scale samples are not analysed using sclerochronological methods yet. These are stored by the County Governor of Innlandet and the Lands museum (https://randsfjordmuseet.no/lands-museum). These scale samples may be used for sclerochronological - and potentially also genetic analysis - in the future (in the context of genetic analyses, stored samples for the period 1966-2003 are also available).

<u>Contact</u>: County Governor of Innlandet

#### Released stocked trout

Several additonal types of data have been collected in connection with the stocking programme. Numbers, body size distribution (as discrete classes), and location of all hatchery smolt releases have been recorded for most years from 1965 onwards, and these data were digitised and aggregated for the period 1965-2018 by Chloé R. Nater and Marlene W. Stubberud. Furthermore, a considerable number of stocked smolt were individually marked upon release as part of experimental studies predominantly between 1970 and 1990. Mark-recapture-recovery data on over 13,000 hatchery reared smolt (including 10,000+ harvest events and around 200 recaptures in the fish ladder) have been separated out of the here described CMRR dataset and are available upon request (note, however, that due to this separation happening at an earlier stage, parts of these data have not been harmonised / error-checked completely).

<u>Contact</u>: Chloé R. Nater

#### Fecundity

A small set of data on female fecundity (number of eggs held by a female of a certain size) has been collected by Chloé R. Nater, Asbjørn Vøllestad, and Yngvild Vindenes during the spawning season in 2017 and 2018. Sample size is small (15 females), but the data may be used to infer fecundity under consideration of typical relationships of female size and fecundity in salmonid fishes (see *Nater et al. 2019a* for use in a population model) and is available upon request.

<u>Contact</u>: Chloé R. Nater

#### Telemetry

The movement of brown trout in River Gudbrandsdalslågen, especially with regards to hydropower dams, river regulation, and habitat fragmentation has been the subject of a variety of telemetry studies over time (see, for example, *Arnekleiv and Kraabøl 1996, Kraabøl et al. 2008, Junge et al. 2013*, *Kraabøl et al. 2015*). Combination of the here described long-term data with telemetry data may provide new opportunities in future studies.

<u>Contact</u>: Refer to corresponding author contact information in the above cited literature.

### Availability of related environmental data

Related long-term environmental monitoring data are available for Lake Mjøsa and its catchment, including data on meteorology, hydrology, river and lake water quality and lake biology (see also *Løvik and Moe 2016*).

- Meteorological data can be downloaded from eklima.met.no, from the monitoring stations Kise (12550) and Toten (11500) [specify periods]
- Hydrological data on river discharge and temperature for River Gudbrandsdalslågen are available from The Norwegian Water Resources and Energy Directorate (www.nve.no) [specify periods]
- Water quality and biological data from with 2-weekly monitoring during the growing season (May-Oct) since 1972 are available from NIVA (Løvik & Moe 2016). These data include nutrients and other physico-chemical data from 6 river stations (main river 1976-2018; other rivers 1986-2018); nutrients, secchi depth, temperature, chlorophyll-a and other physico-chemical variables from 4 lake stations (1972-2018); species-level phytoplankton data from all lake stations (main station Skreia 1972-2016; other stations 1994-2018); and species-level zooplankton and *Mysis* data available from Skreia (1976-2018).
- Satellite data including chl-a, colour, Secchi depth and turbidity from MERIS (2002-2012) and OCLI on-board the Sentinel-3 satellite (2016 onwards), supplemented by data from Landsat satellites are available from NIVA
- Other data on the water quality are available from https://vannmiljo.miljodirektoratet.no/ (by the Norwegian Environment Agency)
- Other data on the catchment and water body properties, ecological status, pressures, river basin management plans etc. are available from http://vann-nett.no (by NVE)

<u>Contact</u>: Jannicke Moe

## Supporting information

Suppl. material 1: The relational SUSTAIN trout database

## Acknowledgements

The project SUSTAIN (Research Council of Norway, contract no. 244647/E10) has funded compilation, harmonisation, checking, and publication of the SUSTAIN trout data, as well as the writing of this data paper.

Collection and maintenance of the data has been financed, carried out, and supported by several institutions and individuals. Opplandskraft DA, who operatea the Hunderfossen dam and powerplant, have been responsible for both the stocking programme (since initialization) and the marking and recapturing of trout in the fish ladder (since 1988). Special thanks go Åse Brenden and Frank Hansen for their outstanding contributions to the operation of the hatchery and stocking programme. The “Merkesentralen” of NINA (Norwegian Institute for Nature Research) has been in charge of managing the system for the reports of tags from outside the fish ladder (especially from harvest). We acknowledge the substantial contribution of the local fishers who have voluntarily reported tags and associated information on their catches for over half a century. The County Governors of Oppland and Hedmark (now merged to “Innlandet”) have both played a vital role in combining and maintaining the data from both the marking and recaptures at Hunderfossen and the reported tags. Data maintenance and cleaning has further been supported by The Norwegian Environment Agency. Prof. Thrond O. Haugen (Norwegian University of Life Sciences) has contributed substantially to earlier efforts of partially cleaning and harmonising the data throughout the last 15 years.

## Author contributions

SJM and CRN share the first authorship. The authors have contributed to this data paper according to the following CRediT statements. Contributions that mainly predate the SUSTAIN project (2015-2019) are indicated by an asterisk.

Conceptualization (of data compilation and data paper): LAV, CRN, SJM

Methodology (relational database): SJM

Software (database and supporting algorithms): SJM, CRN

Validation (of compiled and extracted data): CRN, AR

Formal analysis (of extracted data): CRN

Investigation (collection and provision of hunder trout data): PA*, AR*, OH*, TQ*, LAV

Investigation (sclerochronological analyses of scale samples): PA*, AR*

Resources (materials, samples): PA*, AR*, EL*, OH*, TQ*

Data curation: SJM, CRN, AR*, TQ*, OH*

Writing - original draft: SJM, CRN

Writing - review & editing: CRN, SJM, AR, LAV, EL, OH, TQ, PA Visualization: CRN, SJM, EL, AR

Supervision: LAV

Project administration: LAV, SJM, CRN

Funding acquisition: PA* (monitoring programme); LAV, SJM (SUSTAIN project)

## Supplementary material

**Suppl. material 1: The relational SUSTAIN trout database**

**Authors:** S. Jannicke Moe, Chloé R. Nater et al.

**Data type:** Description of the database

**Brief description:** The file describes the unpublished relational database (SUSTAIN trout database) from which the two publised datasets are extracted; (1) Hunder trout mark-recapture-recovery dataset and (2) Hunder trout growth dataset.

## References

Aass P, Nielsen PS, Brabrand Å (1989) Effects of river regulation on the structure of a fast-growing brown trout (Salmo trutta L.) population. Regulated Rivers: Research & Management 3 (1): 255–266. https://doi.org/10.1002/rrr.3450030125

Aass P (1993) Stocking strategy for the rehabilitation of a regulated brown trout (Salmo Trutta L.) river. Regulated Rivers: Research & Management 8: 135–144. https://doi.org/10.1002/rrr.3450080116

Aass P, Kraabøl M (1999) The exploitation of a migrating brown trout (Salmo trutta L.) population; change of fishing methods due to river regulation. River Research and Applications 15: 211–219.

Aass P, Rustadbakken A, Moe SJ, Lund E, Qvenild T (2017) Life-history data on Hunder brown trout (*Salmo trutta*) from Lake Mjøsa, Norway. Freshwater Metadata Journal1- 11. https://doi.org/10.15504/fmj.2017.25

Arnekleiv JV, Kraabøl M (1996) Migratory behaviour of adult fast-growing brown trout (Salmo trutta, L.) In relation to water flow in a regulated Norwegian river. Regulated Rivers: Research & Management 12 (1): 39–49. https://doi.org/10.1002/(SICI)1099-1646(199601)12:1<39::AID-RRR375>3.0.CO;2-#

Bagenal T (1978) Age and growth. Methods for assessment of fish production in fresh waters. International Biological Programme Handbook no 3. Blackwell Scientific Publishing, Oxford, UK.

Carlin B (1955) Tagging of salmon smolts in the River Lagan. Report Institute of Freshwater Research Drottningholm 36: 57–74.

Clutton-Brock T, Sheldon BC (2010) Individuals and populations: the role of long-term, individual-based studies of animals in ecology and evolutionary biology. Trends in Ecology & Evolution 25 (10): 562–573. https://doi.org/10.1016/j.tree.2010.08.002

Elliott JM (1994) Quantitative ecology and the brown trout. Oxford University Press

Fjeldstad H, Pulg U, Forseth T (2018) Safe two-way migration for salmonids and eel past hydropower structures in Europe: a review and recommendations for best practice solutions. Marine and Freshwater Research 69: 1834–1847. https://doi.org/10.1071/MF18120

Haugen TO, Aass P, Stenseth NC, Vøllestad LA (2008) Changes in selection and evolutionary responses in migratory brown trout following the construction of a fish ladder. Evolutionary Applications 1 (2): 319–335. https://doi.org/10.1111/j.1752-4571.2008.00031.x

Hobæk A, Løvik JE, Rohrlack T, Moe SJ, Grung M, Bennion H, Clarke G, Piliposyan G (2012) Eutrophication, recovery and temperature in Lake Mjøsa: detecting trends with monitoring data and sediment records. Freshwater Biology 57 (10): 1998–2014. https://doi.org/10.1111/j.1365-2427.2012.02832.x

Holtan H (1979) The Lake Mjøsa story. Archiv für Hydrobiologie – Beiheft Ergebnisse der Limnologie 13: 242–258.

Jensen A, Aass P (1995) Migration of a fast-growing population of brown trout (*Salmo trutta L.*) through a fish ladder in relation to waterflow and water temperature. Regulated Rivers: Research & Management 10: 217–228. https://doi.org/10.1002/rrr.3450100216

Junge C, Museth J, Hindar K, Kraabøl M, Vøllestad LA (2013) Assessing the consequences of habitat fragmentation for two migratory salmonid fishes. Aquatic Conservation 24 (3): 297–311. https://doi.org/10.1002/aqc.2391

Kovach R, Muhlfeld C, Al-Chokhachy R, Dunham J, Letcher B, Kershner J (2016) Impacts of climatic variation on trout: a global synthesis and path forward. Reviews in Fish Biology and Fisheries 26 (2): 135–151. https://doi.org/10.1007/s11160-015-9414-x

Kraabøl M (2006) Gytebiologi hos Hunderørret i Gudbrandsdalslågen nedenfor Hunderfossen kraftverk. [Spawning biology of brown trout (*Salmo trutta L.*) in River Gudbrandsdalslågen below Hunderfossen Power Plant.]. NINA report 217, 34 pp. [In Norwegian]. [ISBN 82-426-1777-5]

Kraabøl M, Arnekleiv JV, Museth J (2008) Emigration patterns among trout, *Salmo trutta (L.)*, kelts and smolts through spillways in a hydroelectric dam. Fisheries Management and Ecology 15 (4): 417–423. https://doi.org/10.1111/fme.2008.15.issue-4

Kraabøl M, Dervo BK, Museth J (2015) Downstream migration routes and effects of spillwater release through an ice- and trash spillway on downstream migrating kelts and smolts of the Hunder trout at Hunderfossen Power Station in River Gudbrandsdalslågen. Telemetry studies conducted during autumn 2014 and spring 2015. NINA report 1187[In Norwegian]. URL: http://hdl.handle.net/11250/2388741

Kraabøl M, Museth J (2019) Efficiency of a fishway on brown trout (*Salmo trutta*) spawning populations. Vann 54 (4): 295–311. URL: https://vannforeningen.no/dokumentarkiv/e%EF%AC%83ciency-of-a-%EF%AC%81shway-on-brown-trout-salmotrutta-spawning-populations/

Laikre L, Schwartz M, Waples R, Ryman N (2010) Compromising genetic diversity in the wild: unmonitored large-scale release of plants and animals. Trends in Ecology & Evolution 25 (9): 520–529. https://doi.org/10.1016/j.tree.2010.06.013

Linløkken AN, Johansen W, Wilson R (2014) Genetic structure of brown trout*, Salmo trutta* populations from differently sized tributaries of Lake Mjøsa in south-east Norway. Fisheries Management and Ecology 21 (6): 515–525. https://doi.org/10.1111/fme.12101

Lobón-Cerviá J, Sanz N (2018) Brown trout: Biology, ecology and management. Wiley, Hoboken, NJ, USA,. [ISBN 9781119268352] https://doi.org/10.1002/9781119268352

Løvik JE, Moe SJ (2016) Time series of plankton data from Lake Mjøsa, Norway. Freshwater Metadata Journal 18: 1–9. https://doi.org/10.15504/fmj.2016.18

Lyche Solheim A, Thrane J, Skjelbred B, Økelsrud A, Håll J, Kile MR (2019) Tiltaksorientert overvåking i vannområde Mjøsa. Årsrapport for 2018. [Operational monitoring in water region Mjøsa. Yearly report for 2018]. NIVA report 7373-2019, Oslo, 139 pp. [In Norwegian]. URL: http://hdl.handle.net/11250/2565597 [ISBN 978- 82- 577- 7108-9]

Mills J, Teplitsky C, Arroyo B, Charmantier A, Becker PH, Birkhead T, Bize P, Blumstein D, Bonenfant C, Boutin S, Bushuev A, Cam E, Cockburn A, Côté S, Coulson J, Daunt F, Dingemanse N, Doligez B, Drummond H, Espie RM, Festa-Bianchet M, Frentiu F, Fitzpatrick J, Furness R, Garant D, Gauthier G, Grant P, Griesser M, Gustafsson L, Hansson B, Harris M, Jiguet F, Kjellander P, Korpimäki E, Krebs C, Lens L, Linnell JC, Low M, McAdam A, Margalida A, Merilä J, Møller A, Nakagawa S, Nilsson J, Nisbet IT, van Noordwijk A, Oro D, Pärt T, Pelletier F, Potti J, Pujol B, Réale D, Rockwell R, Ropert-Coudert Y, Roulin A, Sedinger J, Swenson J, Thébaud C, Visser M, Wanless S, Westneat D, Wilson A, Zedrosser A (2015) Archiving Primary Data: Solutions for long-term studies. Trends in Ecology & Evolution 30 (10): 581–589. https://doi.org/10.1016/j.tree.2015.07.006

Muhlfeld C, Dauwalter D, Kovach R, Kershner J, Williams J, Epifanio J (2018) Trout in hot water: A call for global action. Science 360 (6391): 2–867. https://doi.org/10.1126/science.aat8455

Muhlfeld CC, Dauwalter DC, D’Angelo VS, Ferguson A, Giersch JJ, Impson D, Koizumi I, Kovach R, McGinnity P, Schöff mann J, Vøllestad LA, Epifanio J (2019) Global status of trout and char: Conservation challenges in the twentyfirst century. In: Kershner JL, Williams JE, Gresswell RE, Lobón-Cerviá J (Eds) Trout and char of the world. American Fisheries Society, Bethesda, Maryland. [ISBN 978-1-934874-54-7].

Nater C, Rustadbakken A, Ergon T, Langangen Ø, Moe SJ, Vindenes Y, Vøllestad LA, Aass P (2018) Individual heterogeneity and early life conditions shape growth in a freshwater top predator. Ecology 99 (5): 1011–1017. https://doi.org/10.1002/ecy.2178

Nater CR, Stubberud MW, Langangen Ø, Rustadbakken A, Moe SJ, Ergon T, Vøllestad A, Vindenes Y (2019a) A future without stocking? The importance of harvest and river regulation for long-term population viability of migratory salmonids. EcoEvoRxiv (Preprint) https://doi.org/10.32942/osf.io/ds7u2

Nater CR, Vindenes Y, Aass P, Cole D, Langangen Ø, Moe SJ, Rustadbakken A, Turek D, Vøllestad LA, Ergon T (2019b) Estimating cause- and size-specific mortality hazard rates using mark-recapture-recovery data. bioRXiv (Pre-print) https://doi.org/10.1101/544742

Panfili J, De Pontual H, Troadec H, Wrigh PJ (2002) Manual of fish sclerochronology.Ifremer-IRD co-edition, 464 pp.

Piccolo JJ, Norrgård JR, Greenberg LA, Schmitz M, Bergman E (2012) Conservation of endemic landlocked salmonids in regulated rivers: a case-study from Lake Vänern, Sweden. Fish and Fisheries 13 (4): 418–433. https://doi.org/10.1111/j.1467-2979.2011.00437.x

Post JR (2013) Resilient recreational fisheries or prone to collapse? A decade of research on the science and management of recreational fisheries. Fisheries Management and Ecology 20: 99–110. https://doi.org/10.1111/fme.12008

Rustadbakken A, L’Abée-Lund JH, Arnekleiv JV, Kraabøl M (2004) Reproductive migration of brown trout in a small Norwegian river studied by telemetry. Journal of Fish Biology 64 (1): 2–15. https://doi.org/10.1111/j.1095-8649.2004.00275.x

Skaala Ø (1992) Genetic population structure of Norwegian brown trout. Journal of Fish Biology 41 (4): 631–646. https://doi.org/10.1111/j.1095-8649.1992.tb02690.x

Van Leeuwen CA, Dalen K, Museth J, Junge C, Vøllestad LA (2018) Habitat fragmentation has interactive effects on the population genetic diversity and individual behaviour of a freshwater salmonid fi sh. River Research and Applications 34 (1): 60–68. https://doi.org/10.1002/rra.3226

WWF (2018) Living Planet Report - 2018: Aiming Higher. Grooten M, Almond REA (Eds). WWF, Gland, Switzerland.

