## Supplementary material for "Long-term mark-recapture and growth data for large-sized migratory brown trout (*Salmo trutta*) from Lake Mjøsa, Norway": Suppl. material 1: The relational SUSTAIN trout database

### Data sources

The database was compiled from four main data sources (Table 1).

**(1) Individual mark list**: table containing individual-level information on marking, sex, origin, length, weight, release site etc. Obtained from the County Governors of Hedmark and Oppland (2016).

**(2) Mark series list**: table containing ranges of mark numbers with common information on release date, release site, origin (wild or stocked) and stage (smolt or spawner). Obtained from the County Governors of Hedmark and Oppland (2016).

**(3) Recaptures dataset**: table containing all information related to recapture events including date/year, location, equipment, length, weight, and other information. Obtained from "Merkesentralen" at NINA (Norwegian Institute for Nature Research), directly and via the County Governor of Oppland (2016).

**(4) Growth dataset**: table containing all information derived from scales (estimated growth, smolting and spawning). Owned and provided by Per Aass and Atle Rustadbakken, mainly digitized by the County Governor of Hedmark; described by Aass et al. 2017.

**Table S.1.** Description of the main fields of the SUSTAIN trout database (see Figure S.1). The column "Published" indicates which fields are published in the SUSTAIN trout survival dataset. Data souces: Individual = individual mark list, “Series”= Mark series list, “Recaptures” = Recaptures dataset, Growth = Growth dataset.

| Table name | Field name | Data type | Values (examples) | Comment | Main data source | Pub-lished |
|---|---|---|---|---|---|---|
| t_Individual | MarkNo | Text | | Unique code of individual's tag. Key field. | Individual | yes |
| t_Individual | MarkDate | Date/Time | | Date of marking (if available) | Individual | yes |
| t_Individual | MarkYear | Number (integer) | | Year of marking (mandatory) | Individual | yes |
| t_Individual | LengthAtMarking | Number (double) | | Measured length at marking (mm) | Individual | no |
| t_Individual | WeightAtMarking | Number (double) | | Measured weight at marking (g) | Individual | no |
| t_Individual | MarkType | Text | Carlin, Floy | Type of mark | Individual | no |
| t_Individual | Sex | Text | female, male | Individual's sex | Individual | yes |
| t_Individual | Origin | Text | wild, stocked | Individual's origin | Individual; Series | yes |
| t_Individual | ReleaseStage | Text | smolt, spawner | Life-history stage at the time of mark and release | Individual; Series | no |
| t_Individual | ReleaseSite | Text | | Location of relese after marking | Individual; Series | no |
| t_Individual | ReleaseSeason | Text | fall, spring | Only for smolt releases: season of release | Individual; Series | no |
| t_Individual | CaptureNoMax | Number (integer) | | The total number of captures per individual (from t_Capture) | Recaptures | yes |
| t_Capture | CaptureNo | Number (integer) | | Capture number (running number) | Recaptures | yes |
| t_Capture | CaptureDate | Date/Time | | Date of capture (if available) | Recaptures | yes |
| t_Capture | CaptureYear | Number (integer) | | Year of capture (mandatory) | Recaptures | yes |
| t_Capture | LengthAtCapture | Number (double) | | Measured length at capture (mm) | Recaptures | yes |
| t_Capture | WeightAtCapture | Number (double) | | Measured weight at capture (g) | Recaptures | yes |
| t_Capture | AgeAtCapture | Number (integer) | | Estimated age at capture (from scales) | Growth | no |
| t_Capture | AgeMaxRiver | Number (integer) | | Estimated maxiumum age in the river phase (before smolting, from scales) | Growth | no |
| t_Capture | AgeMaxLake | Number (integer) | | Estimated number of years since smolting (from scales) | Growth | no |
| t_Capture | AgeAtSmolting | Number (integer) | | Estimated age at smolting (= AgeMaxRiver, from scales), alternatively from marking lists (for smolt releases) | Growth; Individual | no |
| t_Capture | LengthAtSmolting | Number (double) | | Estimated length at smolting (mm, from scales) | Growth | no |

| | | | | | | |
|---|---|---|---|---|---|---|
| t_Capture | AgeAtMaturation | Number (integer) | | Estimated age at the first spawning event (from scales) | Growth | no |
| t_Capture | LengthAtMaturation | Number (double) | | Estimated length at the first spawning event (from scales) | Growth | no |
| t_Capture | NoOfMigrations | Number (integer) | | Estimated number of spawning events (from scales) | Growth | no |
| t_Capture | HatchYear | Number (integer) | | Estimated year of hatching (age = 0, from scales)) | Growth | no |
| t_Capture | CaptureLocation | Text | | Information on capture location (if available) | Recaptures | no |
| t_Capture | CaptureArea | Text | river trap, river above, river below, river, lake, far away | Standardised values derived from more detailed CaptureLocation | Recaptures | yes |
| t_Capture | CaptureEquipment | Text | net, rod, trap, dead in trap, found dead | Standardised values derived from more detailed information | Recaptures | yes |
| t_Capture | StatusBeforeCapture | Text | alive, dead | Standardised values derived from more detailed information | Recaptures | no |
| t_Capture | StatusAfterCapture | Text | alive, dead | Standardised values derived from more detailed information | Recaptures | no |
| t_Capture | CaptureType | Text | See Table 2 | See Table 2 | Recaptures | yes |
| t_GrowthLake | AgeLake | Number (integer) | | Estimated number of years since smolting | Growth | yes |
| t_GrowthRiver | AgeRiver | Number (integer) | | Estimated number of years since smolting | Growth | yes |
| t_GrowthRiver | LengthRiver | Number (double) | | Estimated length at the given AgeRiver | Growth | no |
| t_GrowthLake | LengthLake | Number (double) | | Estimated length at the given AgeLake | Growth | no |
| t_Spawning | SpawningNo | Number (integer) | | Spawning event number | Growth | no |
| t_Spawning | SpawningYear | Number (integer) | | Estimated year of the spawning event (derived from CaptureYear) | Growth | no |
| t_Spawning | LengthAtSpawning | Number (double) | | Estimated length at the given SpawningNo (mm) | Growth | no |
| t_Spawning | AgeLakeAtSpawning | Number (integer) | | Estimated AgeLake at the given SpawningNo | Growth | no |
| t_Spawning | AgeAtSpawning | Number (integer) | | Estimated total age at the given SpawningNo | Growth | no |
| t_Spawning | SpawningNoMax | Number (integer) | | Estimated total number of spawning events | Growth | no |

**Data compilation and database structure**

The database is organised as 5 main tables (Figure S.1) containing information on individual fish (**t_Individual**), captures (**t_Capture**), growth in the river (**t_GrowthRiver**), growth in the lake (**t_GrowthLake**), and spawning events in the river (**t_Spawning**). A description of the main fields is given in Table 1, while more explanation to the table structure is given below. **t_Individual** is a table summarising all individual-level information that does not change over time. Within this table, each individual is defined by its mark number (**MarkNo**), and this is associated with sex, origin (wild or stocked), and various information from the initial marking event including date (only year in some cases), length, weight, whether the individual was marked as a spawner or as smolt (**ReleaseStage**), *etc*. The total number of captures per individual (**CaptureNoMax**) is derived from the table t_Capture.

The table **t_Capture** contains one or more captures for each individual, and each entry is defined by a unique combination of **MarkNo** and **CaptureNo**. The marking event represents the first capture of an individual and is given the value CaptureNo = 1, regardless of whether the fish were marked as spawners or smolts. **t_Capture** contains all information related to the specific captures, such as capture date, location, type, and equipment, as well as the individual's measured length and weight, and status before and after capture (dead or alive). This table also contains life-history information derived from scales such as age and length at smolting and maturation, the number of spawning migrations (stored in **t_GrowthRiver**, **t_GrowthLake** and **t_Spawning**). Such information does, in principle, belong to the individual level as it should not change throughout life. However, this information is derived from scales, and the estimates from scales taken at different captures of the same individual may. Therefore, this information has been stored at the capture level.

**Table S.2.** Definition of the field CaptureType.

| Capture Area | StatusBefore Capture | StatusAfter Capture | Other criteria | Capture Type |
|---|---|---|---|---|
| trap | alive | alive | ReleaseStage = smolt AND CaptureNo = 1 | smolt release |
| trap | alive | alive | ReleaseStage ≠ smolt OR CaptureNo ≠ 1 | trap alive |
| trap | alive | dead | | trap dead |
| river or lake | alive | alive | | catch release |
| river or lake | alive | dead | | harvest |
| river or lake | dead | dead | | found dead |

The complete information on growth derived from the scale analysis is stored in two tables representing the river phase of life (i.e. from birth to smolting; **t_GrowthRiver**) and the lake phase of life (post-smolting and including later spawning runs; **t_GrowthLake**). The reason for estimating lake growth separately is that scales are sometimes lost and replaced by new scales, e.g. after the fish has migrated to the lake. The replacement scales cannot always represent the complete river phase but can still be used to estimate the growth in the lake phase.

**t_GrowthRiver** contains estimated individual records of age and length in each year from the year of smolting (clearly visible on the scale) back to the first year (**AgeRiver** = 1). Likewise, **t_GrowthLake** contains individual records of age and length in each year from the sampling (= capture) year back to the first year in the lake (**AgeLake**= 1). The estimated lake age and river age at capture is stored in t_Capture as **AgeLakeMax** and **AgeRiverMax**, respectively.

The estimated **AgeAtSmolting** corresponds to **AgeRiverMax** when this is available. (For fish marked as smolt rather than as spawners, the **AgeAtSmolting** is known). In cases where information from the river phase is intact, the **AgeAtCapture** is the sum of **AgeRiverMax** and **AgeLakeMax**.

All information on individual spawning schedules derived from scale analysis are stored in **t_Spawning.** This table contains the estimated year, age (or **AgeLake**) and length for all spawning events of each individual. The **AgeAtMaturation** and **LengthAtMaturation** field in **t_Capture** correspond to the estimated age and length at the first spawning event here.

**Database contents: summary**

The complete **SUSTAIN trout database** contains and links mark-recapture data and individual-level data on growth trajectories and spawning schedules of the Hunder trout over 50 years. The mark-recapture data contains altogether 43,249 capture records from all years 1966-2017 (Table 3). Of these, 27,773 are individual marking events from the years 1966-2015, representing either the first capture of mature trout in the ladder (14,624 individuals) or the release of tagged smolt (13,149 individuals). The remaining capture records are recaptures by different means (as defined by **CaptureType**): 2,190 recaptures in the trap (378 of which resulted in death); 12,905 harvest reports (8,976 in the lake,3796 in the river and 133 in other or unknown locations); 105 reports of fish found dead; and 20 reports of catch-and-release fishing. The individual-level data on growth trajectories contains 29,940 size-at-age records for 5,115 individuals. The individual-level data on spawning schedules contains 7,189 spawning records of 5,007 individuals. All these individuals have been captured in the Hunderfossen fish ladder during the period 1966-2005 (Aass et al., 2017) and are also contained in the mark-recapture data.

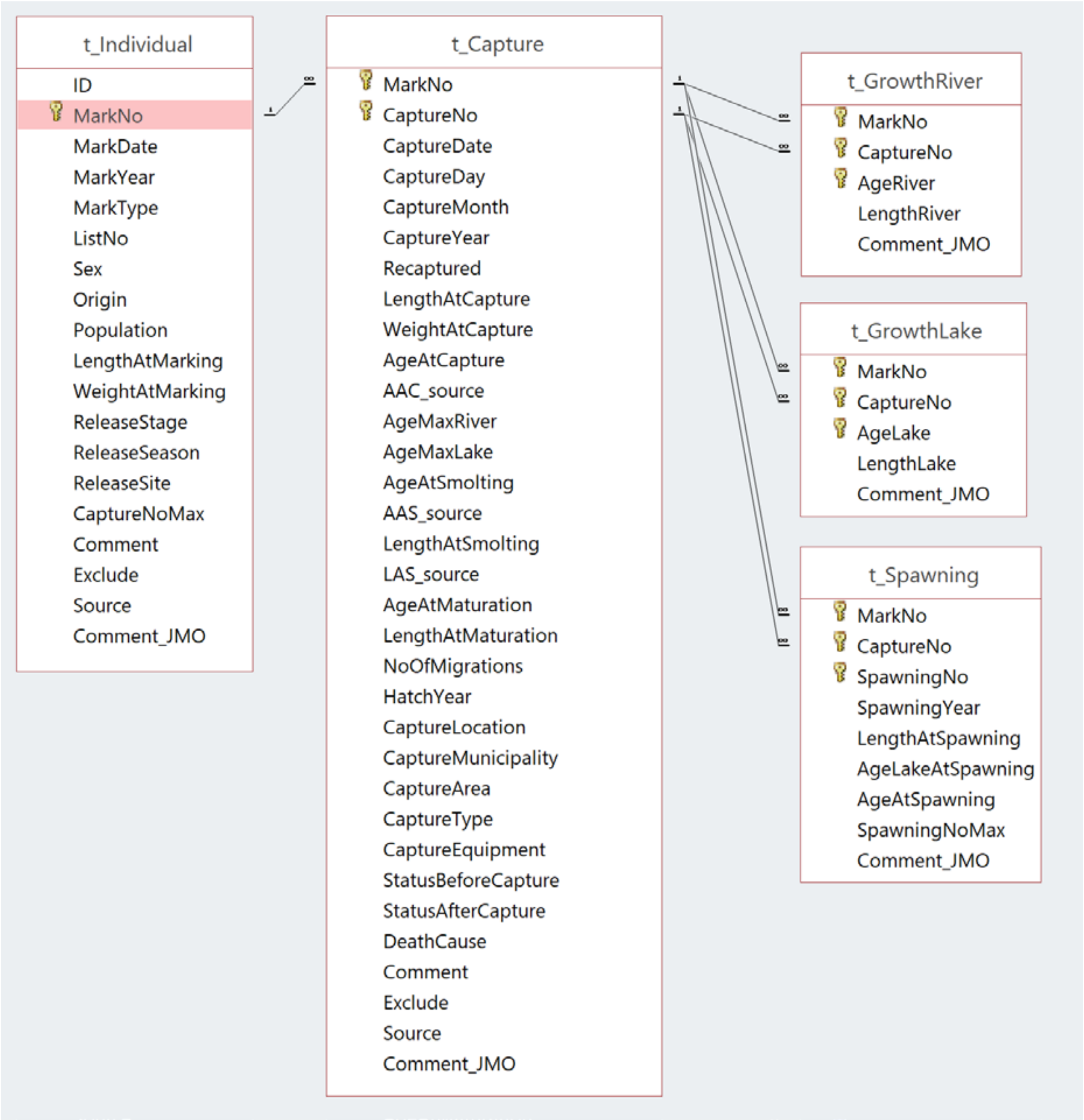


**Figure S.1.** Illustration of the data model for the relational SUSTAIN trout database. The connecting lines illustrate one-to-many relationship from t_Individual to t_Capture, and from t_Capture to the three tables t_GrowthRiver, t_GrowthLake and t_Spawning. The key symbol illustrates the single or composite key field(s) identifying the unique records of each table. After Løvik & Moe (2016).

#### Extraction of datasets for data analysis and publication

The majority of the mark-recapture data from the **SUSTAIN trout database** has been extracted and organised into a single flat table for analysing survival of mature adult trout (Nater et al. 2019a, b). This dataset conditions on the adult trout having passed the Hunderfossen fish ladder on a spawning migration at least once and was extracted from the database using certain quality criteria.

The growth data were extracted and organised into a single flat file for analysing the effects of individual heterogeneity and early life conditions on the growth of trout (Nater et al. 2018).

Both datasets are published with this data paper.
